# A frequent variant in the Japanese population determines quasi-Mendelian inheritance of rare retinal ciliopathy

**DOI:** 10.1101/257634

**Authors:** Konstantinos Nikopoulos, Katarina Cisarova, Mathieu Quinodoz, Hanna Koskiniemi-Kuending, Noriko Miyake, Pietro Farinelli, Atta Ur Rehman, Muhammad Imran Khan, Andrea Prunotto, Masato Akiyama, Yoichiro Kamatani, Chikashi Terao, Fuyuki Miya, Yasuhiro Ikeda, Shinji Ueno, Nobuo Fuse, Akira Murakami, Yuko Wada, Hiroko Terasaki, Koh-Hei Sonoda, Tatsuro Ishibashi, Michiaki Kubo, Frans P. M. Cremers, Zoltán Kutalik, Naomichi Matsumoto, Koji M. Nishiguchi, Toru Nakazawa, Carlo Rivolta

## Abstract

Hereditary retinal degenerations (HRDs) are Mendelian diseases characterized by progressive blindness and caused by ultra-rare mutations. In a genomic screen of 331 unrelated Japanese patients, we identify a disruptive *Alu* insertion and a nonsense variant (p.Arg1933*) in the ciliary gene *RP1*, neither of which are rare alleles in Japan. p.Arg1933* is almost polymorphic (frequency = 0.6%, amongst 12,000 individuals), does not cause disease in homozygosis or heterozygosis, and yet is significantly enriched in HRD patients (frequency = 2.1%, i.e. a 3.5-fold enrichment; p-value = 9.2×10^−5^). Familial co-segregation and association analyses show that p.Arg1933* can act as a Mendelian mutation, *in trans* with the *Alu* insertion, but might also cause disease in association with two alleles in the *EYS* gene in a non-Mendelian pattern of heredity. Our results suggest that rare conditions such as HRDs can be paradoxically determined by relatively common variants, following a quasi-Mendelian model linking monogenic and complex inheritance.

## INTRODUCTION

Together with intellectual disabilities, hereditary retinal degenerations (HRDs, comprising retinitis pigmentosa and allied diseases) represent a group of conditions for which both genetic and allelic heterogeneity is the highest in humans^1,2^. To date, almost 300 genes and thousands of mutations have been identified as causative of HRD, and the detection of novel disease genes and variants continues at a steady pace (RetNet database: https://sph.uth.edu/retnet/). Considering that the overall prevalence of HRDs does not exceed 1 in 2,000 individuals, the average contribution of any given HRD gene to the disease is incredibly small. Similarly, apart from two DNA variants that appear to be relatively frequent in the general population and determine a specific form of the disease (p.Asn1868Ile and p.Gly863Ala in *ABCA4*)^3,4^, most mutations are so rare that they are seldom detected in more than one pedigree, worldwide. In addition, although HRDs affect people from the five continents, their specific allelic assortment seems to be population-specific^5,6^. For instance, similar to other islanders or groups of people that have experienced relative historical isolation, Japanese carry certain alleles, including pathogenic ones, which are not found elsewhere in the world^7^. Furthermore, lack of significant reduction in fitness before the reproductive age, associated with such an elevated heterogeneity, have led to the consequence that the number of recessive mutations that are detected heterozygously in the general, unaffected population is remarkably high and may affect up to one person in two^8^.

Despite this extraordinary variability and abundance of mutations, HRD is almost invariantly inherited as a monogenic, Mendelian trait, for which the presence of only one (dominant) or two (recessive) mutations in the same gene, genome-wide, is at the same time a necessary and sufficient condition for pathogenicity^9^. At the other end of the spectrum of ocular conditions having a genetic component lies age-related macular degeneration (AMD), another retinal disease affecting people aged 50 and over. AMD is a *bona fide* complex disease with a relatively high prevalence (1 in 13 individuals), favored by the presence of polymorphic SNPs, highly-penetrant rare variants, and environmental factors^10^. Between these two pillars of inheritance, there is an intermediate zone, consisting in a few examples for which extremely rare mutations in more than one gene are associated with Bardet-Biedl syndrome, a retinal ciliopathy displaying sometimes digenic triallelic inheritance^11–13^.

*RP1* is one of the several HRD genes identified to date, and one of the few causing disease by more than one Mendelian pattern of inheritance. Originally described as linked to autosomal dominant retinitis pigmentosa (adRP)^14–16^, a subtype of HRD, it was later shown to be associated with a recessive form of the same disease (arRP)^17^. To date, at least 60 mutations have been reported in *RP1*, most of which cluster within its last exon (exon 4), cumulatively accounting approximately for 5.5% and up to 4.5% of all adRP and arRP cases, respectively^18,19^. However, some DNA variants at the far 3’ end of the gene, including nonsense variants, appear not to cause disease, at least not according to a dominant or recessive pattern of inheritance^20,21^. *RP1* encodes a multi-modular protein of 2,156 amino acids, which is a member of the doublecortin family and is present in the ciliary axoneme of both rods and cones, the light-sensing neurons of the retina^22,23^. Mutations in *RP1* thus determine visual loss as a consequence of a ciliopathic phenotype affecting these specialized cell types.

Following the screen of a large set of Japanese HRD patients, we identify three mutations in the *RP1* gene: an unusual mobile *Alu* element insertion in exon 4, a novel frameshift mutation, and a nonsense variant in the far 3’ part of the coding sequence. While the first two variants behave as classical recessive Mendelian alleles, p.Arg1933* appears to cause disease according to a more complex pattern of inheritance. When present *in trans* with respect to the *Alu* insertion, it acts as a Mendelian mutation. Furthermore, despite being enriched in patients vs. controls, p.Arg1933* is completely benign in homozygosis or in heterozygosis. By performing an association test between 28 HRD patients, heterozygous carriers of this nonsense allele, and 3,554 controls, we find that p.Arg1933* may be pathogenic not only as a Mendelian allele, but also in association with variants elsewhere in the genome, and in particular with two DNA changes in another ciliary gene, *EYS*.

## RESULTS

### An *Alu* insertion in *RP1* is a prevalent cause of HRD in Japan

In the framework of a screening effort of 331 unrelated Japanese patients, we identified a novel, unusual mutation by Whole-Genome Sequencing, consisting in the insertion of a mobile *Alu* element in exon 4 of the *RP1* gene (m1, or NM_006269.1:c.4052_4053ins328/p.Tyr1352Alafs*9) in a female individual from a recessive HRD family. This insertion caused the disruption of the reading frame by introducing 328 additional nucleotides, including a premature termination codon in the canonical *RP1* coding sequence. The mother of the proband was heterozygous for this variant and the proband’s affected brother was also a homozygote, in support of the notion that this was indeed a recessive HRD mutation (Fig. 1a). Targeted screening for this *Alu* insertion in the remaining 330 patients (all forms of HRDs, isolate or recessive cases, not genetically pre-screened) as well as in 524 Japanese controls, available for direct testing of m1, identified 15 other affected and unrelated individuals and one heterozygous control carrying this insertion. In total, six patients were homozygous for the mutation (12 alleles), which co-segregated with the disease as a classical Mendelian, recessive allele, whenever this could be tested, while 10 carried it heterozygously. Altogether, these findings indicate that this *Alu* insertion is not only clearly pathogenic, but it is also a rather prevalent cause of retinal degeneration within the Japanese islands at the level of a single allele (1.8% of all HRD Japanese patients), possibly second only to the most frequent mutation so far identified in this country, i.e. NM_001142800.1:c.4957dup in *EYS*^24–26^.

**Figure 1.**
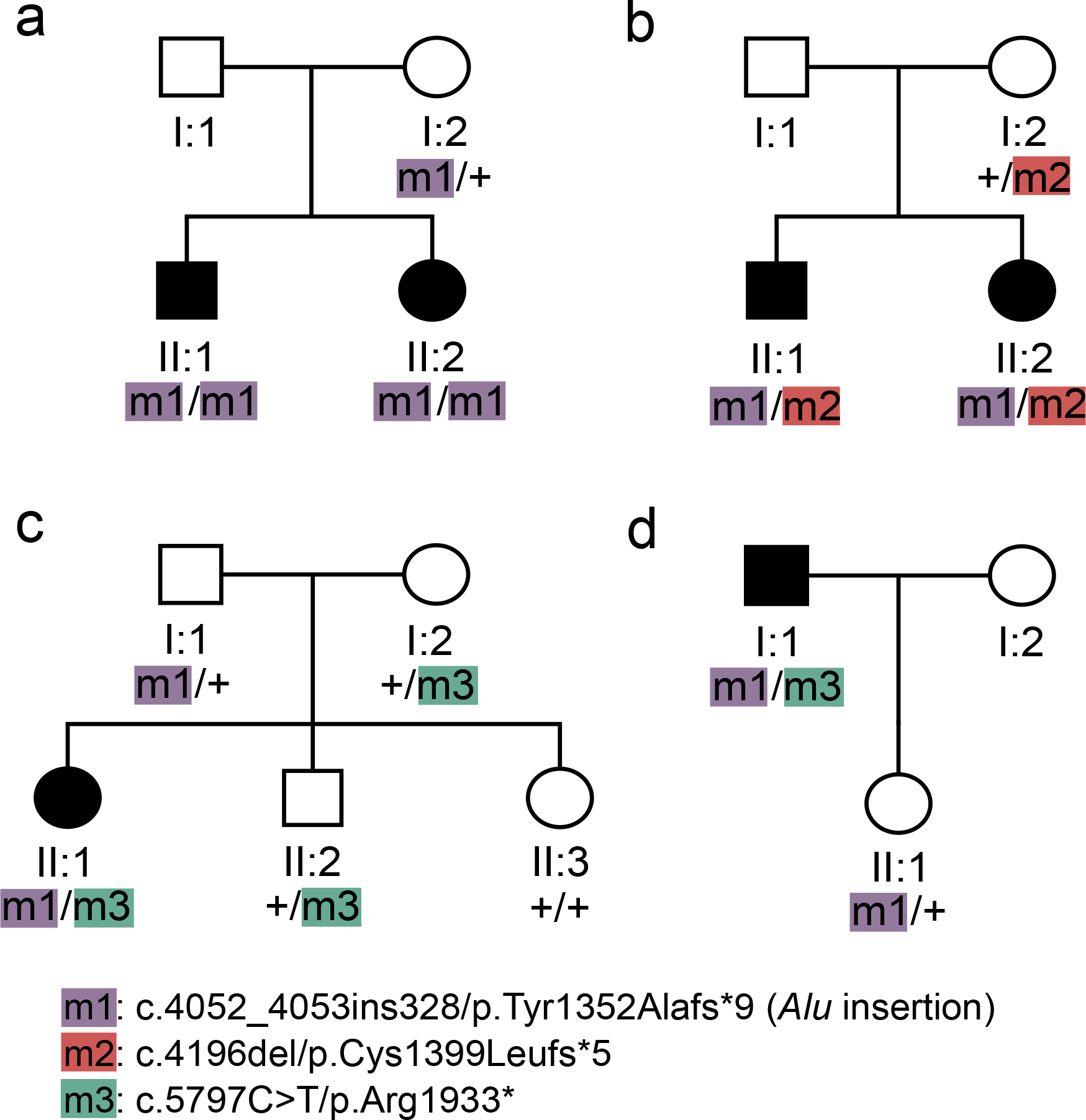
Segregation analysis of the *RP1* mutations found in this study. Pedigrees of representative families are shown.

Remarkably, 6 of the 10 individuals who carried the *Alu* insertion heterozygously were in fact compound heterozygotes for either of two other changes in *RP1*: a novel frameshift mutation (c.4196del/p.Cys1399Leufs*5, m2, two unrelated individuals) and a nonsense variant c.5797C>T/p.Arg1933* (m3, four unrelated individuals) that was previously identified in the general population and is present in dbSNP as entry # rs118031911 (Supplementary Table 1). Again, both variants, detected by direct Sanger sequencing, co-segregated with the disease in relevant families, according to an autosomal recessive pattern of inheritance (Fig. 1bcd).

**Table 1.** Summary of the results of the association study. The top 10 hits from this test are shown. OR, odds ratio; CI, confidence interval; *, *p*-values retaining significance following a Bonferroni correction for multiple testing, over 178 variants and for α = 0.05.

| Gene | Variant | Frequency<br>in cases | Frequency<br>in controls | OR | 95% CI (OR) | *p*-value |
| --- | --- | --- | --- | --- | --- | --- |
| *EYS* | NM_001142800.1:c.2528G>A; p.Gly843Glu | 0.125 | 0.017 | 8.23 | 3.08-18.78 | 5.6E-05* |
| *EYS* | NM_001142800.1:c.8805C>A; p.Tyr2935* | 0.054 | 0.002 | 33.30 | 5.86-128.76 | 1.9E-04* |
| *EYS* | NM_001142800.1:c.4957del; p.Ser1653Valfs*26 | 0.054 | 0.004 | 13.85 | 2.62-46.83 | 1.9E-03 |
| *USH2A* | NM_206933.2:c.15355C>T; p.Arg5119Trp | 0.036 | 0.002 | 20.15 | 2.16-92.44 | 5.9E-03 |
| *USH2A* | NM_206933.2:c.2802T>G; p.Cys934Trp | 0.036 | 0.003 | 14.55 | 1.60-63.22 | 0.010 |
| *PDZD7* | NM_024895.4:c.1267G>A; p.Ala423Thr | 0.036 | 0.003 | 13.38 | 1.48-57.70 | 0.012 |
| *EYS* | NM_001142800.1:c.7394C>G; p.Thr2465Ser | 0.089 | 0.029 | 3.28 | 1.01-8.30 | 0.024 |
| *CC2D2A* | NM_001080522.2:c.501G>T; p.Lys167Asn | 0.036 | 0.004 | 8.44 | 0.96-34.65 | 0.027 |
| *RPGRIP1L* | NM_015272.4:c.171G>T; p.Leu57Phe | 0.054 | 0.011 | 4.90 | 0.96-15.63 | 0.028 |
| *BBIP1* | NM_001243783.2:c.112T>C; p.Ser38Pro | 0.036 | 0.005 | 7.70 | 0.87-31.35 | 0.032 |

### m3 is enriched in patients, but does cause HRD *per se*

Frameshift c.4196del/p.Cys1399Leufs*5 (m2) was absent from 3,480 Japanese control chromosomes and was reported in the gnomAD database^27^ to have an allele frequency of 5.44×10^−5^ in East Asia, indicating that this DNA variant is a very rare allele, as it is the case for most HRD mutations.

In contrast, the rs118031911/T allele (m3), despite being virtually absent in many world populations, was found to be relatively frequent in East Asians (Supplementary Figure 1), and probably too frequent to be a Mendelian allele for HRD, according to the Hardy-Weinberg model. In particular, our direct screening of 12,379 Japanese individuals with no retinal degeneration showed the presence of rs118031911/T in 145 subjects, 142 heterozygotes and 3 homozygotes (148 alleles), validating the notion that this DNA variant is in fact almost polymorphic in Japan (allele frequency = 0.6%). All these subjects were examined by fundoscopy and, in addition, we evaluated clinically one of the three homozygotes (the only one who could be re-assessed, in agreement with our Institutional Review Board protocol) by a very thorough ophthalmological examination. At age 28 y.o., she had no visual symptoms and displayed no ocular abnormalities: she had normal visual acuity (20/20 in both eyes), intact visual field (Goldmann perimetry), and no evidence of retinal degeneration through slit lamp examination and fundoscopy. Furthermore, optical coherence tomography imaging, used to assess detailed retinal structures, showed no sign of retinal thinning and electroretinogram, a test allowing objective detection of minimal retinal dysfunction even in the absence of subjective symptoms, showed normal responses. Finally, absence of late-onset HRD, who could have escaped detection in a 28 y.o. individual, was confirmed by the assessment of the fundi of the other two rs118031911/T homozygotes, who displayed no signs of retinal degeneration at ages of 78 and 79 years, respectively. Overall, both population based-data and direct clinical assessments confirm that rs118031911/T does not cause HRD *per se*, in heterozygosis or in homozygosis.

However, specific screening for the rs118031911/T allele in the same cohort of 331 Japanese HRD patients mentioned above led to the identification of 10 additional heterozygotes (14 alleles in total) showing that its frequency in HRD patients was 2.1% (14 alleles out of 662) (Supplementary Figure 1). The 3.5-fold enrichment of rs118031911/T in patients vs. controls (148 alleles out of 24,758 = 0.6%) was highly significant [*p*-value = 9.2×10^−5^, threshold = 0.05/*N*, *N*=1, by Fisher’s exact test (24,610:148 vs. 648:14)], indicating that this relatively common variant has in fact an effect on retinal health. WES analysis of these 10 patients detected no mutations in HRD genes that could explain their phenotype, according to a Mendelian fashion of inheritance. Considering that rs118031911/T introduces a nonsense codon in the *RP1* open reading frame and was found *in trans* with respect to the *Alu* insertion in some patients, it is not unlikely that it could represent a hypomorphic variant contributing to the mutational load of genes involved in retinal homeostasis. In other words, despite being benign when considered as a Mendelian allele (monoallelically or biallelically), rs118031911/T could exert a pathogenic function in conjunction with DNA changes in other HRD genes, according to an oligogenic pattern of inheritance that was previously modeled for hereditary ciliopathies^28–30^.

### A non-Mendelian pattern of inheritance for m3

We tested this hypothesis by assessing for enrichment of nonsynonymous, rare and low-frequency variants (minor allele frequency between 0.1% and 5%, according to published literature^31–34^; further details in Methods) in the 10 patients mentioned above as well as 18 additional patients with the same genotype (heterozygous for rs118031911/T, with no other recognized mutations in *RP1* and no mutations in HRD genes that could explain their phenotype), identified following a targeted screening of 713 Japanese HRD cases from another internal cohort (Supplementary Table 1). Specifically, we performed an association test between these 28 individuals and 3,554 Japanese controls from the 3.5KJPN database^35^ by considering all 228 *bona fide* HRD genes^36^ from the RetNet database (Supplementary Table 2) that could produce multiallelic inheritance of HRD in m3 heterozygotes, in line with previous protocols involving similar analyses^37–42^. Cryptic relatedness among patients as well as the presence of additional, undetected *RP1* mutations *in trans* with respect to rs118031911/T were excluded prior to performing the test (Supplementary Tables 3 and 4). The association analysis identified two variants that were significantly enriched in patients vs. controls (Table 1, Fig. 2). Interestingly, although they were not in linkage disequilibrium, both variants belonged to the gene *EYS* (NM_001142800.1:c.2528G>A;p.Gly843Glu and c.8805C>A;p.Tyr2935*, *p*-values = 5.6×10^−5^ and 1.9×10^−4^, respectively; threshold = 2.8×10^−4^ = 0.05/*N*, where *N*=178, by Fisher’s exact test), possibly highlighting a mechanism of pathogenesis directly involving the proteins EYS and RP1. Indeed, a third DNA change within *EYS* (c.4957del;p.Ser1653Valfs*26) ranked 3^rd^ in the list of associated variants, even if its *p*-value did not reach statistical significance after Bonferroni correction. Furthermore, we repeated the same analysis by considering not only variants from RetNet sequences, but from the whole human exome, in both cohorts. In support of the robustness of the data obtained, the two significant hits detected above were also the two top hits detected exome-wide, although obviously no variant reached the threshold for multiple corrections on this overly large number of additional tests. Altogether, these results indicate that the rs118031911/T nonsense could act in concert with at least two DNA changes (and possibly with more) to determine a pathological phenotype in a non-Mendelian fashion.

**Figure 2.**
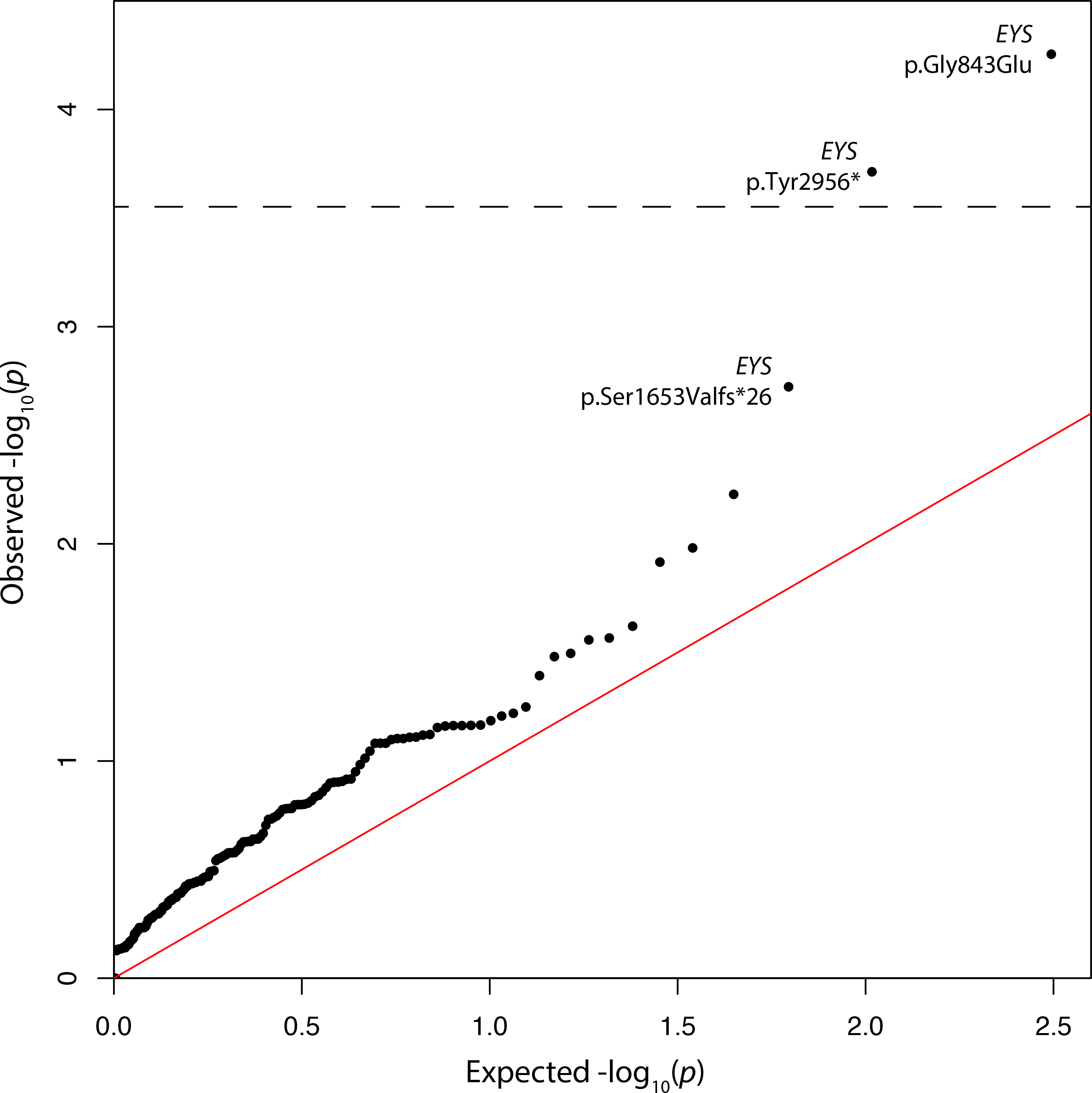
Results of the association study. Quantile-quantile (Q-Q) plot of rare / low-frequency non-synonymous variants in HRD genes in 28 patients heterozygous for rs118031911/T vs. 3,554 Japanese controls. The significance threshold is indicated by the dotted line. Nomenclature for variants in the *EYS* gene are relative to entry NM_001142800.1.

Based on previous data on Bardet-Biedl syndrome^11^, we tested a digenic diallelic vs. triallelic mode of action for rs118031911/T on HRD, by comparing the frequency of this variant in patients for whom the molecular causes of retinal degenerations were identified (i.e. solved cases) vs. unsolved HRD cases vs. controls. As expected, a comparison of unsolved vs. controls showed significance, as reported above, whereas solved vs. controls did not show any significant enrichment for rs118031911/T (6 rs118031911/T variants over 722 alleles for solved vs. 148 over 24,758 for controls, *p*-value = 0.46, OR = 1.4, CI = 0.50-3.14, by Fisher’s exact test). Comparison of solved vs. unsolved HRD cases showed borderline non-significant enrichment for rs118031911/T in unsolved cases (*p*-value = 0.07, OR = 2.30, CI = 0.94-6.76, by Fisher’s exact test), possibly indicating that either well-defined triallelism does not take place for this variant or, simply, that we did not have enough power to detect it.

Similarly, we tested whether the significant hits detected in *EYS* in patients carrying rs118031911/T heterozygously were not the trivial consequence of *EYS* being commonly involved in RP in Japan^24^, a feature potentially leading to false positive results following the comparison of patients (enriched in mutations in *EYS*, in general, regardless of the *RP1* genotype) vs. controls (depleted for these same variants). Comparison of genotypes in rs118031911/T heterozygous patients vs. unsolved HRD patients not carrying rs118031911/T showed that both the p.Gly843Glu and p.Tyr2935* variants were enriched in the former group of individuals [*p*-value = 0.039, OR = 2.30 (19.6% vs. 9.57%), CI = 0.73-13.3, for p.Gly843Glu and *p*-value = 0.054, OR = 3.88 (6.52% vs. 1.76%), CI = 0.96-4.96 for p.Tyr2935*, by Fisher’s exact test], confirming that the increased presence of these DNA changes in *EYS* was truly due to their association with the rs118031911/T genotype.

## DISCUSSION

The extreme genetic heterogeneity of retinal degenerations, together with the elevated number of pathogenic and hypomorphic changes in HRD genes that are detected in the unaffected population, have evoked the theoretical possibility that non-Mendelian, oligogenic inheritance could be responsible for these conditions^9^. Digenic heredity has been clearly demonstrated for specific combinations of mutations^43–45^ in particular pedigrees or in individual cases, including digenic triallelic transmission of Bardet-Biedl syndrome^11,46^. For these patients, the presence of two (diallelic) or three (triallelic) mutations at two different loci (digenism) causes disease, presumably by compromising the overall function of gene products that belong to the same complex or are part of the same biochemical pathway. This model seems to be particularly true for genes encoding for proteins that form or play a role within the cell primary cilium, according to the paradigm of mutational load put forward by N. Katsanis and coworkers^47^. In these instances, accumulation of rare variants (which individually may have a little effect) in multiple ciliary genes can produce a pathological phenotype that is connected to ciliary function and result in a ciliopathy, including retinal ciliopathies^48–50^.

Intriguingly, despite our association test was not limited to ciliary genes, both our significant hits lied within a ciliary gene, *EYS*. In primates, the EYS protein has been shown to physically co-localize with RP1 in the ciliary axoneme of photoreceptors and is thought to play a role in the structural organization and maintenance of these cells’ apical part, the outer segment (OS)^51^. This functional role is further supported by studies in zebrafish, where *EYS* knockouts show progressive retinal degeneration due to mis-localization of specific OS proteins and the disruption of F-actin filaments^52,53^, a key component not only for the integrity, but also for the morphogenesis of the OS^54^. In a similar fashion, targeted disruption of the *RP1* gene in mice leads to defects of the OS, because of the incorrect stacking of its discs^55^. The co-localization of RP1 and EYS, as well as their common role in the homeostasis of the OS, strongly indicates that they may have synergic functions and that pathogenesis could occur in a digenic fashion.

In this work we show that two specific *RP1* alleles are responsible for a relatively large number of Mendelian HRD cases in Japan. Interestingly, none of these two changes is a rare allele at all, compared to the average frequencies of classical HRD mutations. The first, the c.4052_4053ins328/p.Tyr1352Alafs*9 *Alu* element insertion in *RP1*, seems to be the second most common HRD recessive mutation described so far in Japan, and its frequency may even be underestimated, since insertional events of mobile elements are difficult to detect by conventional screening techniques. The second variant, c.5797C>T/p.Arg1933* or rs118031911/T, is even more frequent, and by far more interesting. Despite introducing a premature stop codon in the *RP1* open reading frame, this DNA change is almost polymorphic in East Asia and does not cause disease either in heterozygous or homozygous carriers. However, this same change may act as pathogenic allele in a Mendelian fashion (with another *RP1* mutation *in trans*), or in association with rare variants in at least another gene, according to a non-Mendelian, possibly oligogenic pattern of inheritance. Although we currently ignore the molecular mechanisms leading to this unusual model of pathogenicity, it is probably the consequence of an increased global mutational load with threshold effect, determined by the accumulation of variants with different pathogenic potential. The presence of one or of two rs118031911/T alleles likely produces a load that is below this pathological threshold, while the co-occurrence of extra variants could result in the crossing of such a limit for normal retinal homeostasis. This hypothesis is supported by the evidence that rs118031911/T is pathogenic in conjunction with a very severe mutation, i.e. the insertion of an *Alu* element in *RP1*’s exon 4 mentioned above, which completely ablates the open reading frame of the gene. We term this model of inheritance quasi-Mendelian, to define the differential behavior (Mendelian or non-Mendelian) that specific alleles may have with respect to different genotypes at the same locus or elsewhere in the genome.

In conclusion, it seems that, at least for *RP1*-associated HRD, disorders displaying a Mendelian pattern of inheritance may also genetically behave like multigenic conditions, for which both polymorphic (having a low effect) and rare (having a rather high effect) variants can determine pathogenesis (Figure 3). Although our study clearly needs to be replicated, possibly in other East Asian cohorts, the low prevalence of HRD and the even lower percentage of HRD patients carrying rs118031911/T prevents us at the moment to increase the power of our analysis and to propose more defined models of pathogenicity, since the identification of the 28 heterozygotes used in this study corresponds roughly to the screening of 5-10 million Japanese individuals. However, this work provides a clear proof of concept that a non-negligible proportion of HRDs can be caused by inheritance mechanisms that transcend the Mendelian model, to be investigated in detail by future, very large-scale and population-specific sequencing endeavors, such as for instance the 100,000 genomes project^56^. Furthermore, our findings suggest that oligogenic heredity of human diseases (and perhaps of other characters) may not be limited to a low number of cases with hyper-rare conditions, as shown up to now^38,39,57,58^, but could extend to more frequent phenotypes and represent a bridge between monogenic and complex inheritance.

**Figure 3.**
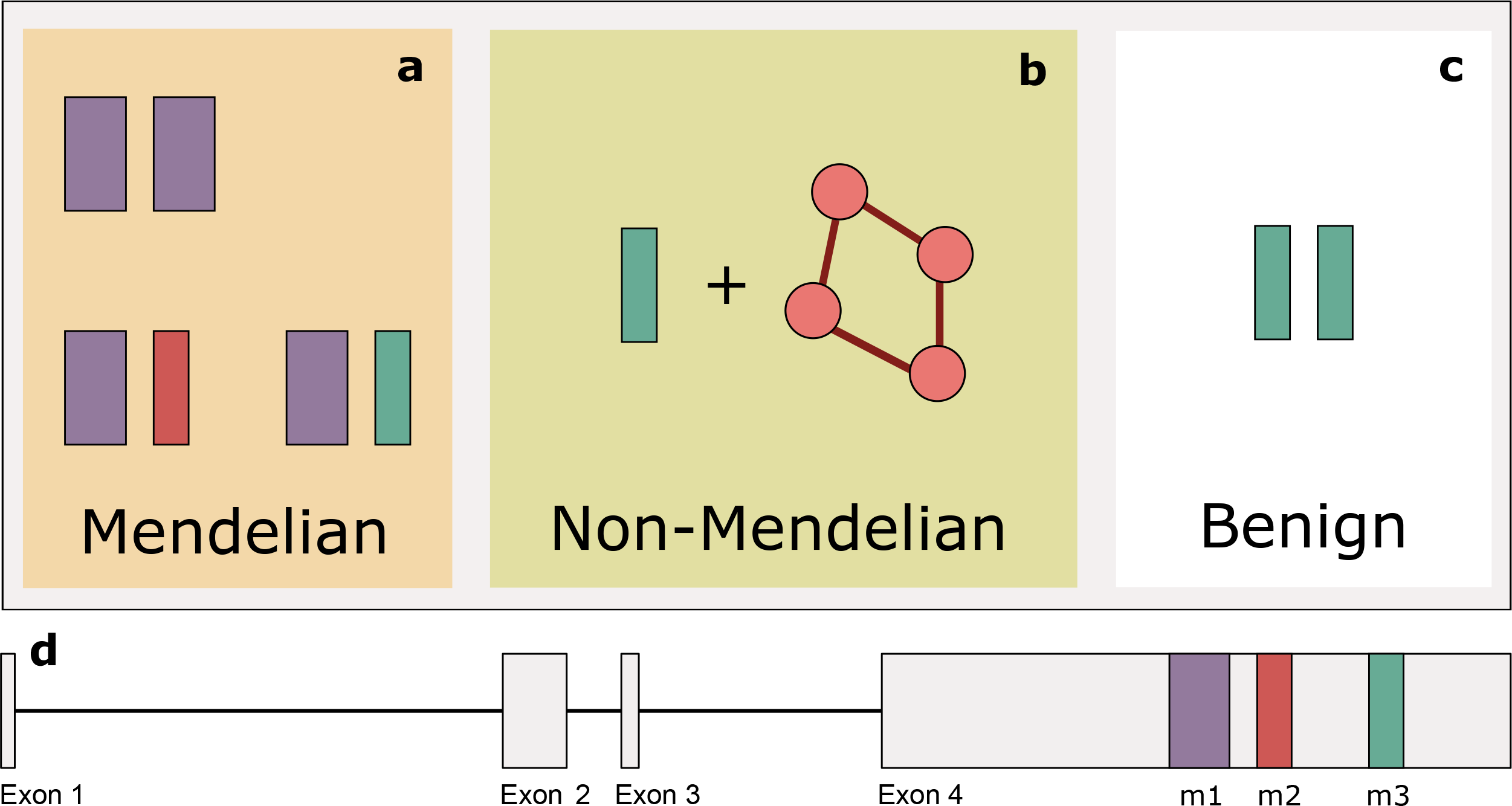
Schematic representation of the inheritance pattern of the identified mutations in *RP1*, highlighting the concept of rs118031911/T-mediated quasi-Mendelian inheritance of HRDs. (a) Any combination of the *RP1 Alu* element insertion (m1, or c.4052_4053ins328/p.Tyr1352Alafs*9), in a homozygous state or in a compound heterozygous combination with m2 (c.4196del/p.Cys1399Leufs*5) or rs118031911/T (m3, or c.5797C>T/p.Arg1933*) results in autosomal recessive inheritance of the disease. (b) Combinations of the hypomorphic m3 allele with additional hypomorphs and/or heterozygous recessive alleles in other genes results in disease following a non-Mendelian pattern, whereas (c) homozygosis for m3 has no pathological consequences. (d) Structure of *RP1*: exons are represented by boxes, connected by solid lines (introns). The relative positions of m1, m2, and m3 are also indicated.

## METHODS

### Subjects

The study was initiated following the approval by the Institutional Review Boards of our respective Institutions. All subjects provided written informed consent, and the study was conducted in adherence with the Declaration of Helsinki.

Tohoku University School of Medicine, Kyushu University School of Medicine, and Nagoya University School of Medicine, all based in Japan, were the centers where all Japanese patients with HRD were recruited. HRD was diagnosed clinically after excluding possible secondary causes of retinal degeneration such as toxicity and uveitis. Final diagnosis required the presence of reduced electroretinogram (ERG) responses, visual field loss, and funduscopic abnormalities consistent with retinal degeneration (retinal vascular narrowing and abnormalities of the retinal pigment epithelium etc.) symmetrically in both eyes.

Genotypes from individuals without HRD were collected from both published and unpublished databases [the BioBank Japan Project (N=12,379), the ToMMo Japanese Reference Panel Project (3.5KJPN release, N=3,554)^35^, the Tohoku University School of Medicine (N=95), the Yokohama City University Graduate School of Medicine (N=429)] or were obtained experimentally by direct genotyping of genomic DNA, with standard molecular biology techniques.

Summary phenotypes of carriers of m1, m2, or m3 mutations are listed in Supplementary Table 1.

### Whole genome sequencing and analysis

Genome sequencing of the first index patient was performed using the sequencing platform by Complete Genomics^59^. Sequence reads were mapped to the human reference genome (NCBI build 37) and variants were called genome-wide. These included: single-nucleotide variants (SNVs), copy-number variations (CNVs), as well as structural variations (SVs) such as *Alu* element insertions and/or chromosomal rearrangements. Data were extracted from MasterVar files and other relevant matrices by ad hoc Perl, bash, and R scripts, available upon request. Assessment of pathogenic variants was performed as previously described^60^.

### Screening for the *Alu* element insertion

In order to screen for the presence of the *Alu* element in exon 4 of *RP1* distinct pair of primers were designed (forward: 5′-AGGCTTGTTTCCTAGGAGAGGT-3′, reverse: 5′-TTCTGCTTCTTTTTCACTTAGGC-3′) using the CLCbio Genomics Workbench (Qiagen, Hilden, Germany).

PCR amplification was performed in a 20 μl total volume containing 20 ng genomic DNA, 1x GoTaq buffer, 0.5 mM dNTPs, 10 μM of each primer, and 2 units (5 U/μl) of GoTaq polymerase (Promega, Madison, Wisconsin). PCR products were separated following agarose gel electrophoresis. PCR products displaying abnormal size profiles were purified (ExoSAP-IT, USB, Cleveland Ohio) and a sequencing reaction was performed in a total volume of 5 μl using 1 μl primer 3.3 μM, 0.5 μl BigDye Terminator v1.1, and 1 μl of the provided Buffer (Applied Biosystems, Foster City, California) Big Dye terminator cycle sequencing kit on an ABI 3130xl Genetic Analyzer (Applied Biosystems).

For this screening, we used 524 controls from the Tohoku University School of Medicine (N=95) and the Yokohama City University Graduate School of Medicine (N=429).

### Whole exome sequencing and analysis

Paired-end DNA sequencing libraries of 28 individuals were generated using Aglilent SureSelect Human All ExonV6 kit (Agilent Technologies, CA, USA) by Novogene Co., Ltd., Hong Kong. One microgram of genomic DNA per sample was fragmented into 180-280bp fragments by hydrodynamic shearing (Covaris, Massachusetts, USA). After the reparation of the 3’ and 5’ ends and the adenylation of the 3’ ends, paired-end adaptors were ligated to the DNA fragments. DNA fragments with ligated adaptors on both ends were enriched by PCR. PCR products were further purified using the AMPure XP system (Beckman Coulter, Beverly, USA) and quantified using the Agilent high sensitivity DNA assay on the Agilent Bioanalyzer 2100 system. Captured DNA fragments were sequenced on an Illumina NovaSeq 6000 platform (Illumina, San Diego, California).

#### Mapping, variant calling and annotation

Raw sequence files were assessed, trimmed, and mapped to the human genome reference sequence (UCSC hg19) using Novoalign V3.08.02 (Novocraft, Selangor, Malaysia). Variants were called jointly by GATK 3.8^61^ and annotation was performed using the combination of Annovar^62^ and an in-house developed annotation pipeline^63^. The nomenclature of all variants studied was validated by using VariantValidator^64^.

#### Filtering

All single nucleotide variants were further filtered to obtain only high-quality variants. Briefly, quality control was carried out using the following parameters: (1) remove individual calls if Depth (DP) < 8 or GenotypeQuality (GQ) < 20, (2) exclude variants if the average GQ value ≤ 35, (3) exclude variants if call-rate value ≤ 0.9, (4) keep only variants with no deviation from Hardy-Weinberg equilibrium (p>0.05 after Bonferroni correction), (5) keep variants passing GATK VQSR (VQSRTranche of 90.0), (6) final hard filtering step with Quality by Depth (QD) ≥ 2, FisherStrand (FS) ≤ 60, RMSMappingQuality (MQ) ≥ 40, MappingQualityRankSumTest (MQRankSum) ≥ −12.5, ReadPosRankSumTest (ReadPosRankSum) ≥ −8, StrandOddsRatio (SOR) ≤ 3 and ExcessHet ≤ 20.

### Association study on rare and low-frequency variants in RetNet genes

An association test was performed on rare and low-frequency non-synonymous variants from a curated RetNet list (N=228 genes, Suppl. Table 2) to test possible association of variants in the 28 patients carrying rs118031911/T heterozygously compared to 3,554 controls from the ToMMo database^35^. More specifically, variants were retained if they had a frequency comprised between 0.1% and 5% in controls, with the exclusion of rs118031911/T itself, according to published methods^31–34^. This resulted in the selection of 178 variants in 84 different genes, which were used to test association in rs118031911/T carriers vs. controls by Fisher’s exact test with an experiment-wide Bonferroni-corrected threshold of 2.81×10^−4^, for α = 0.05 (0.05 / 178 = 2.81×10^−4^). P-values and odds ratios were obtained by the fisher.test function with default parameters in R (v3.5.1) and the Q-Q plot (Fig. 2) was obtained by using the qqman package^65^.

### Relatedness analyses

PLINK (v1.90b5)^66^ was used to compute PI_HAT values between all pairs of the 28 rs118031911/T carriers using calls from whole exome sequencing. This analysis showed no relatedness, with PI_HAT values between 0.00 and 0.07 (threshold for relatedness = 0.2)^67^, for all 378 possible pairwise combinations (Supplementary Table 3).

### SNP genotyping of the *RP1* locus

To detect haplotypes *in trans* with respect to rs118031911/T, we genotyped 23 SNPs encompassing the *RP1* locus over ~260 kb, by standard techniques.

## ACKNOWLEDGEMENTS

This work was supported by the following agencies. The Swiss National Science Foundation (grant #176097, to C.R.); the Rotterdamse Stichting Blindenbelangen, the Stichting voor Ooglijders and the Nelly Reef fund (to M.I.K. and F.P.M.C.); the Japan Agency for Medical Research and Development (grant #17ek0109213h0001) and the Japan Society for the Promotion of Science (grant #16K11315, to K.M.N.); Research on Measures for Intractable Diseases, Comprehensive Research on Disability Health and Welfare, the Strategic Research Program for Brain Science, and the Initiative on Rare and Undiagnosed Diseases (all to N.Ma.); the Japanese Agency for Medical Research and Development, Grants-in-Aid for Scientific Research (to N.Ma. and N.Mi.); the BioBank Japan Project, by the Japanese Ministry of Education, Culture, Sport, Science and Technology and the Japanese Agency for Medical Research and Development (to M.K.); the PhD Fellowship in Life Sciences (Faculty of Biology and Medicine, University of Lausanne), to MQ. The authors would also like to warmly thank Dr. Matthew Robinson for his expert advice on this work.

