## Supplementary Figure 1 for "A frequent variant in the Japanese population determines quasi-Mendelian inheritance of rare retinal ciliopathy"

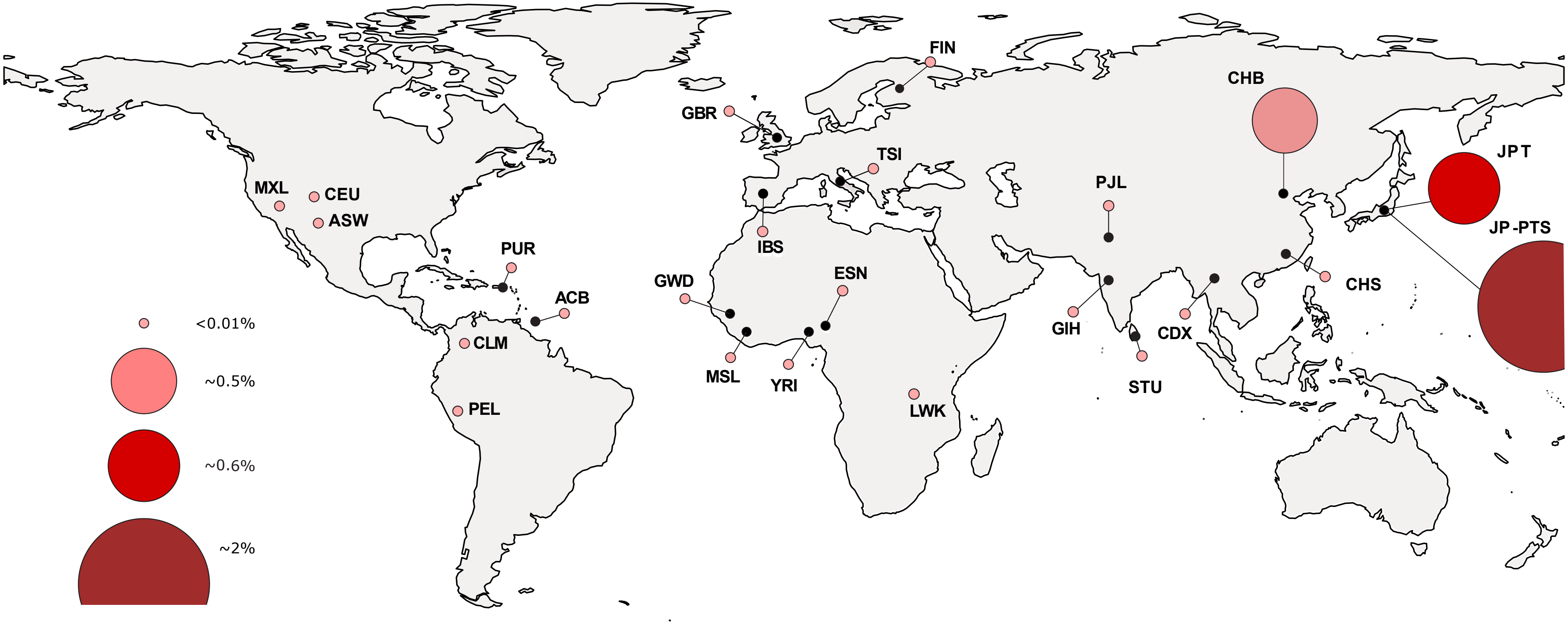


**Supplementary Figure 1.** Frequency of the rs118031911/T allele across different world populations. ACB, African Caribbeans in Barbados; ASW, Americans of African Ancestry in South West USA; BEB, Bengali from Bangladesh; CDX, Chinese Dai in Xishuangbanna, China; CEU, Utah Residents (CEPH) with Northern and Western European Ancestry; CHB, Han Chinese in Bejing, China; CHS, Southern Han Chinese; CLM, Colombians from Medellin, Colombia; ESN, Esan in Nigeria; FIN, Finnish in Finland; GBR, British in England and Scotland; GIH, Gujarati Indian from Houston, Texas; GWD, Gambian in Western Divisions in the Gambia; IBS, Iberian Population in Spain; JPT, Japanese in Tokyo, Japan; JP-PTS, Japanese HRD patients from our set; LWK, Luhya in Webuye, Kenya; MSL, Mende in Sierra Leone; MXL, Mexican Ancestry from Los Angeles USA; PEL, Peruvians from Lima, Peru; PJL, Punjabi from Lahore, Pakistan; PUR, Puerto Ricans from Puerto Rico; STU, Sri Lankan Tamil from the UK; TSI, Toscani in Italia; YRI, Yoruba in Ibadan, Nigeria. This figure was generated by using the R software.
