## Supplementary Table 1 for "A frequent variant in the Japanese population determines quasi-Mendelian inheritance of rare retinal ciliopathy"

**Supplementary Table 1.** Clinical data for patients heterozygous for m3 or with biallelic m1, m2, or m3 mutations. MD, macular dystrophy; CD, cone dystrophy; CRD cone-rod dystrophy; RP, retinitis pigmentosa.

| Patient ID | Diagnosis | Age at examination | Gender | Age of symptoms onset | Genotype |
|---|---|---|---|---|---|
| M15 | MD | 73 | M | 53 | m3/+ |
| M9 | CD | 84 | F | 60 | m3/+ |
| R(O)40 | RP | 37 | F | elementary school | m3/+ |
| R21 | RP | 63 | M | unknown | m3/+ |
| 221 | RP | 67 | M | ~30 | m3/+ |
| R(O)44 | RP | 30 | F | 18 | m3/+ |
| OPH-610 | RP | 48 | F | 20 | m3/+ |
| OPH-302 | RP central type | 37 | F | childhood | m3/+ |
| Q-116 | RP | 60 | F | childhood | m3/+ |
| OPH-179 | RP | 78 | M | 20 | m3/+ |
| OPH-285 | RP | 55 | F | 40 | m3/+ |
| OPH-419 | RP | 44 | F | 18 | m3/+ |
| OPH-424 | RP | 84 | M | 10 | m3/+ |
| OPH-733 | RP | 68 | M | 50 | m3/+ |
| OPH-794 | RP | 50 | M | 40 | m3/+ |
| OPH-812 | RP | 80 | M | 26 | m3/+ |
| OPH-814 | CRD | 51 | F | 20 | m3/+ |
| OPH-884 | RP | 58 | F | 40 | m3/+ |
| OPH-967 | RP | 37 | M | 25 | m3/+ |
| R170 | RP | 31 | M | 31 | m3/+ |
| R204 | RP | 47 | F | 4 | m3/+ |
| YWC101 | RP | 41 | M | 8 | m3/+ |
| YWC102 | RP | 44 | M | 30 | m3/+ |
| YWC107 | RP | 73 | F | 65 | m3/+ |
| OPH-553 | RP | 60 | M | 40 | m3/+ |
| YWC100 | RP | 42 | F | 20 | m3/+ |
| YWC193 | RP | 69 | F | 20 | m3/+ |
| YWC6 | RP | 42 | F | 35 | m3/+ |
| R(O)39 | RP | 29 | F | elementary school | m1/m1 |
| R(O)48 | RP | 12 | F | 12 | m1/m1 |
| R(O)70 | RP | 34 | F | elementary school | m1/m1 |
| RN36 | RP | 32 | M | 6 | m1/m1 |
| OPH-635 | RP | 46 | M | 12 | m1/m1 |
| 08_20 | RP | 54 | F | 30s | m1/m1 |
| 201 | RP | 28 | F | 5 | m1/m2 |
| 209 | RP | 26 | F | childhood | m1/m2 |
| C1 | MD | 41 | F | 25 | m1/m3 |
| M5 | MD | 55 | M | 44 | m1/m3 |
| M7 | MD | 43 | F | 36 | m1/m3 |
| OPH-280 | RP | 61 | F | 50 | m1/m3 |
