## Supplementary Table 2 for "A frequent variant in the Japanese population determines quasi-Mendelian inheritance of rare retinal ciliopathy"

**Supplementary Table 2.** HRD genes (from  RetNet) used for the association test (N=228).

| | | | | | |
|---|---|---|---|---|---|
| ABCA4 | CEP164 | GPR179 | MKKS | PRPF3 | SNRNP200 |
| ABCC6 | CEP250 | GRK1 | MKS1 | PRPF31 | SPATA7 |
| ABHD12 | CEP290 | GRM6 | MVK | PRPF4 | SPP2 |
| ACBD5 | CEP78 | GUCA1A | MYO7A | PRPF6 | TIMM8A |
| ADAM9 | CERKL | GUCA1B | NBAS | PRPF8 | TIMP3 |
| ADAMTS18 | CFAP410 | GUCY2D | NDP | PRPH2 | TMEM216 |
| ADGRA3 | CHM | HARS | NEK2 | PRPS1 | TMEM237 |
| ADGRV1 | CIB2 | HGSNAT | NMNAT1 | RAB28 | TOPORS |
| AGBL5 | CLN3 | HK1 | NPHP1 | RAX2 | TRIM32 |
| AHI1 | CLRN1 | HMX1 | NPHP3 | RBP3 | TRNT1 |
| AHR | CLUAP1 | IDH3A | NPHP4 | RBP4 | TRPM1 |
| AIPL1 | CNGA1 | IDH3B | NR2E3 | RCBTB1 | TTC8 |
| ALMS1 | CNGA3 | IFT140 | NRL | RD3 | TTLL5 |
| ARHGEF18 | CNGB1 | IFT172 | NYX | RDH11 | TTPA |
| ARL2BP | CNGB3 | IFT27 | OAT | RDH12 | TUB |
| ARL3 | CNNM4 | IFT81 | OFD1 | RDH5 | TUBGCP4 |
| ARL6 | CRB1 | IMPDH1 | OPN1LW | REEP6 | TUBGCP6 |
| ARSG | CRX | IMPG1 | OPN1MW | RGR | TULP1 |
| ASRGL1 | CSPP1 | IMPG2 | PANK2 | RGS9 | USH1C |
| ATF6 | CTNNA1 | INPP5E | PCARE | RGS9BP | USH1G |
| BBIP1 | CWC27 | INVS | PCDH15 | RHO | USH2A |
| BBS1 | CYP4V2 | IQCB1 | PCYT1A | RIMS1 | WDPCP |
| BBS10 | DHDDS | ITM2B | PDE6A | RLBP1 | WDR19 |
| BBS12 | DHX38 | JAG1 | PDE6B | ROM1 | WFS1 |
| BBS2 | DRAM2 | KCNJ13 | PDE6C | RP1 | WHRN |
| BBS4 | DTHD1 | KCNV2 | PDE6G | RP1L1 | ZNF408 |
| BBS5 | EFEMP1 | KIAA1549 | PDE6H | RP2 | ZNF423 |
| BBS7 | ELOVL4 | KIF11 | PDZD7 | RP9 | ZNF513 |
| BBS9 | EMC1 | KIZ | PEX1 | RPE65 | |
| BEST1 | EXOSC2 | KLHL7 | PEX2 | RPGR | |
| C1QTNF5 | EYS | LAMA1 | PEX7 | RPGRIP1 | |
| C8orf37 | FAM161A | LCA5 | PHYH | RPGRIP1L | |
| CABP4 | FLVCR1 | LRAT | PITPNM3 | RS1 | |
| CACNA1F | FZD4 | LRIT3 | PLK4 | SAG | |
| CACNA2D4 | GDF6 | LRP5 | PNPLA6 | SAMD11 | |
| CC2D2A | GNAT1 | LZTFL1 | POC1B | SDCCAG8 | |
| CCT2 | GNAT2 | MAK | POC5 | SEMA4A | |
| CDH23 | GNB3 | MERTK | POMGNT1 | SLC24A1 | |
| CDH3 | GNPTG | MFRP | PRCD | SLC25A46 | |
| CDHR1 | GPR125 | MFSD8 | PROM1 | SLC7A14 | |
