## Supplementary Table 4 for "A frequent variant in the Japanese population determines quasi-Mendelian inheritance of rare retinal ciliopathy"

**Supplementary Table 4.** Haplotype surrounding rs118031911/T (in bold) across ~260 kb, in 11 randomly-selected patients out of the 28 p.Arg1933* heterozygotes considered for the association test. Positions are given with respect to the hg19 genome build.

| Chromosome | Position | Change | OPH-967 | YWC101 | YWC102 | YWC107 | OPH-179 | OPH-812 | OPH-794 | OPH-884 | OPH-814 | OPH-419 | OPH-424 |
|---|---|---|---|---|---|---|---|---|---|---|---|---|---|
| 8 | 55505788 | A>G | G | G | G | A | G | G | G | G | G | G | G |
| 8 | 55506499 | T>G | T | T | T | T | T | T | T | T | T | G | G |
| 8 | 55513201 | A>G | A | G | A | A | A | A | G | A | A | G | A |
| 8 | 55522880 | G>A | G | A | G | G | G | G | A | G | G | G | G |
| 8 | 55523753 | A>G | A | A | A | A | A | A | A | A | A | G | A |
| 8 | 55529612 | T>C | C | T | C | T | T | C | T | T | C | T | C |
| 8 | 55539057 | C>T | T | C | T | C | C | T | C | C | T | C | C |
| 8 | 55539395 | T>A | T | T | T | T | T | T | T | T | T | A | T |
| 8 | 55541450 | C>T | T | C | T | C | C | T | C | C | T | C | C |
| 8 | 55541513 | A>G | G | A | G | A | A | G | A | A | G | A | A |
| **8** | **55542239** | **C>T** | **T** | **T** | **T** | **T** | **T** | **T** | **T** | **T** | **T** | **T** | **T** |
| 8 | 55542540 | C>T | C | C | C | C | C | C | C | C | C | T | C |
| 8 | 55552310 | T>C | T | T | T | T | T | T | T | T | T | C | T |
| 8 | 55629852 | A>G | A | G | A | G | G | A | G | G | A | A | G |
| 8 | 55632762 | G>A | G | A | G | A | A | G | A | A | G | G | A |
| 8 | 55678538 | T>G | T | G | T | G | G | T | G | G | T | T | G |
| 8 | 55688171 | G>A | G | A | G | A | A | G | A | A | G | G | G |
| 8 | 55692780 | A>G | A | G | A | G | G | A | G | G | A | A | G |
| 8 | 55693133 | C>T | C | C | C | C | C | C | C | C | C | T | C |
| 8 | 55735722 | C>T | C | T | C | T | C | T | T | T | C | C | C |
| 8 | 55756256 | G>A | G | G | G | G | G | G | G | G | G | A | G |
| 8 | 55761124 | A>G | A | G | A | G | G | A | G | G | A | G | G |
| 8 | 55763773 | G>A | G | A | G | A | A | G | A | A | G | G | G |
